# Ultrastructural features of presynaptic active zones and dense core vesicles of olfactory projection neuron boutons in *Drosophila melanogaster*

**DOI:** 10.1101/2021.01.12.426300

**Authors:** Kai Yang, Tong Liu, He Liu, Zengru Di, Ke Zhang

**Author notes:** Corresponding author (K. Z.).

## Abstract

In *Drosophila melanogaster*, olfactory projection neurons (PNs) convey odor information from the peripheral olfactory center to higher brain regions. The anatomical and physiological properties of PNs have been well characterized at the cellular and circuit level. The ultrastructural features of PNs remain unknown however, particularly with respect to presynaptic active zones (PAZs) and dense core vesicles (DCVs). In the current study, membrane-labeled electron microscopy was used to volume-reconstruct 89 PN axonal boutons and identify the internal PAZs and DCVs. Based on ultrastructural parameters, these PN boutons could be classified into three morphological distinct subtypes. Interestingly, the distributions of PAZs and DCVs were distinct within these three subtypes. DCVs were enriched in membrane labeled GH146-positive boutons, suggesting that GH146-positive PNs release both neurotransmitters and neuromodulators. The study identified the detailed distributions of PAZs and DCVs in PN boutons and indicates that neuromodulators mediated by DCVs may play an important role in PNs for olfactory processing.

## Introduction

A well as classical neurotransmitters, neuromodulators play essential roles in regulating physiological functions and behaviors in the central nervous system [1–4]. How a neuron releases neurotransmitters or neuromodulators is a critical question in the field of neuroscience. Previous studies have shown that before release, neurotransmitters are stored in small clear core vesicles that drift or dock at presynaptic active zones (PAZs), and neuromodulators are stored in large dense core vesicles (DCVs) that drift in the cytoplasm [5, 6]. Some neuropeptides reportedly serve as cotransmitters that modulate synaptic plasticity [7–10], indicating that DCVs may be morphologically close to PAZs in axonal terminals, but questions about DCV distribution patterns and the relationships between DCVs and PAZs remain to be answered. The olfactory projection neurons (PNs) in adult fruit flies (*Drosophila melanogaster*) provide a convenient model with which to investigate these questions.

In *D. melanogaster’s* olfactory circuit, PNs relay odor information from the primary olfactory center, the antenna lobe, to two brain regions, the mushroom body (MB) and the lateral horn (LH), regulating adaptive and innate behaviors respectively[11]. The calyx is the input region of the MB, where PNs transform dense-coding odor information to sparse-coding odor information in MB Kenyon Cells (KCs) by forming a distinct structure called the microglomerulus [12, 13]. Multiple mechanisms reportedly underlie the coding transformation between PNs and KCs, including thresholding, input integration, sparse synaptic connection, and developmental plasticity [14–17]. There is no direct evidence that neuromodulators mediate this process, but recent transcriptome data [18] and electron microscopy (EM) results [13] suggest that PNs may release neuropeptide as a neuromodulator at their axonal boutons, as well as two types of neurotransmitters, acetylcholine and γ-aminobutyric acid [19, 20].

In the current study, 89 PN boutons in a reference region in the MB calyx were obtained via horse radish peroxidase (HRP)-labeled EM volume reconstruction. We demonstrate that: (1) Based on ultrastructural morphological parameters, these PN boutons can be classified into three types; complex bouton, unilobed bouton, and simple bouton; (2)PAZ number was linearly correlated with bouton volume, whereas there was no strong correlation between DCV number and bouton volume;(3) There were significantly more PAZs and DCVs in complex bouton than in unilobed and simple boutons; (4) GH-146-positive PNs (HRP positive) had significantly larger numbers of DCVs than GH-146-negative (HRP-negative) PNs. In summary, the study analyzed the distribution features of PAZs and DCVs in PN boutons, and shed light on neuromodulator function in olfactory information processing and plasticity in *D. melanogaster*.

## Materials and methods

### *D. melanogaster* strains

The y^1^ w^1118^; GH146-GAL4/CyO strain (Bloomington Drosophila Stock Center: 30026) was used to drive UAS-CD2:HRP/CyO (Bloomington Drosophila Stock Center: 8763).

### EM sample preparation

EM sample preparation was performed as previously described [21], with the modifications described below. Brains from 4-day-old adult female *D. melanogaster* were dissected in 2.5% glutaraldehyde in 0.1 M phosphate buffer (PB; pH 7.4) on a cold plate at 4°C, then fixed overnight at 4°C. After three 10-minute washes with 0.1 M TriHCl buffer (pH 7.4) at 4°C, brains were stained with 1 mg/mL DAB in 0.1 M TriHCl (pH 7.4) for 20 minutes at 4°C, then transferred to room temperature for 15 minutes. After three 10-minute washes with 0.1 M PB, brains were fixed with 1% OsO_4_ in 0.1 M PB for 2 hours at 4°C. After three 10-minute washes with double-distilled water, brains were stained with 2% uranyl acetate overnight at 4°C. Dehydration and infiltration with resin were performed in accordance with standard protocols [22].

### EM data acquisition

EM samples were trimmed and oriented vertically with the MB calyx on the top. Serial EM images were then acquired via focused ion beam scanning EM (Helios Nanolab 600i). The EM sample was imaged in x and y with 5-nm pixels in a small region of 20 × 20 μm in the center of the MB calyx. A focused beam of 30-kV gallium atoms was used to mill the EM sample every 40 nm to acquire a raw dataset of 658 slices.

### Statistical methods

The skeleton length, segment number, volume, and surface area were calculated via Fiji-TrakEM2. PAZs and DCVs were annotated and exported via the customized dissector function in Fiji-TrakEM2. The Kolmogorov–Smirnov test was used to assess the normality of all data distributions. The unpaired *t*-test was used to compare two groups, and one-way analysis of variance was used to compare more than two groups. Spearman’s correlation coefficients were calculated for skeleton length and segment number, and Pearson’s correlation coefficients were calculated for volume and DCV number, and PAZ number. Different models were used to obtain a high cluster quality, Akaike information criteria for morphology clustering, and Bayesian information criteria for PAZ-DCV clustering [23, 24].

## Results

### PN bouton volume reconstruction in a reference region of the MB calyx

PNs connect to the MB in the calyx region to process olfactory information (Figure 1A). The calyx exhibits a round shape from a posterior view under EM, and it is surrounded by KC cell bodies and PN processes (Figure 1B). In the MB calyx a microglomerulus contains large presynaptic PN axonal boutons, numerous tiny postsynaptic dendritic claws, and some other MB extrinsic neurons [19]. In the current study, focused ion beam scanning EM was used to collect a serial EM dataset in a volume of 20 × 20 × 25 μm in the center of the MB calyx to quantitively analyze the properties of PN axonal boutons. The resolution in the x–y axis was 5 nm, and the resolution in the z axis was 40 nm (Figure 1C).

**Figure 1:**
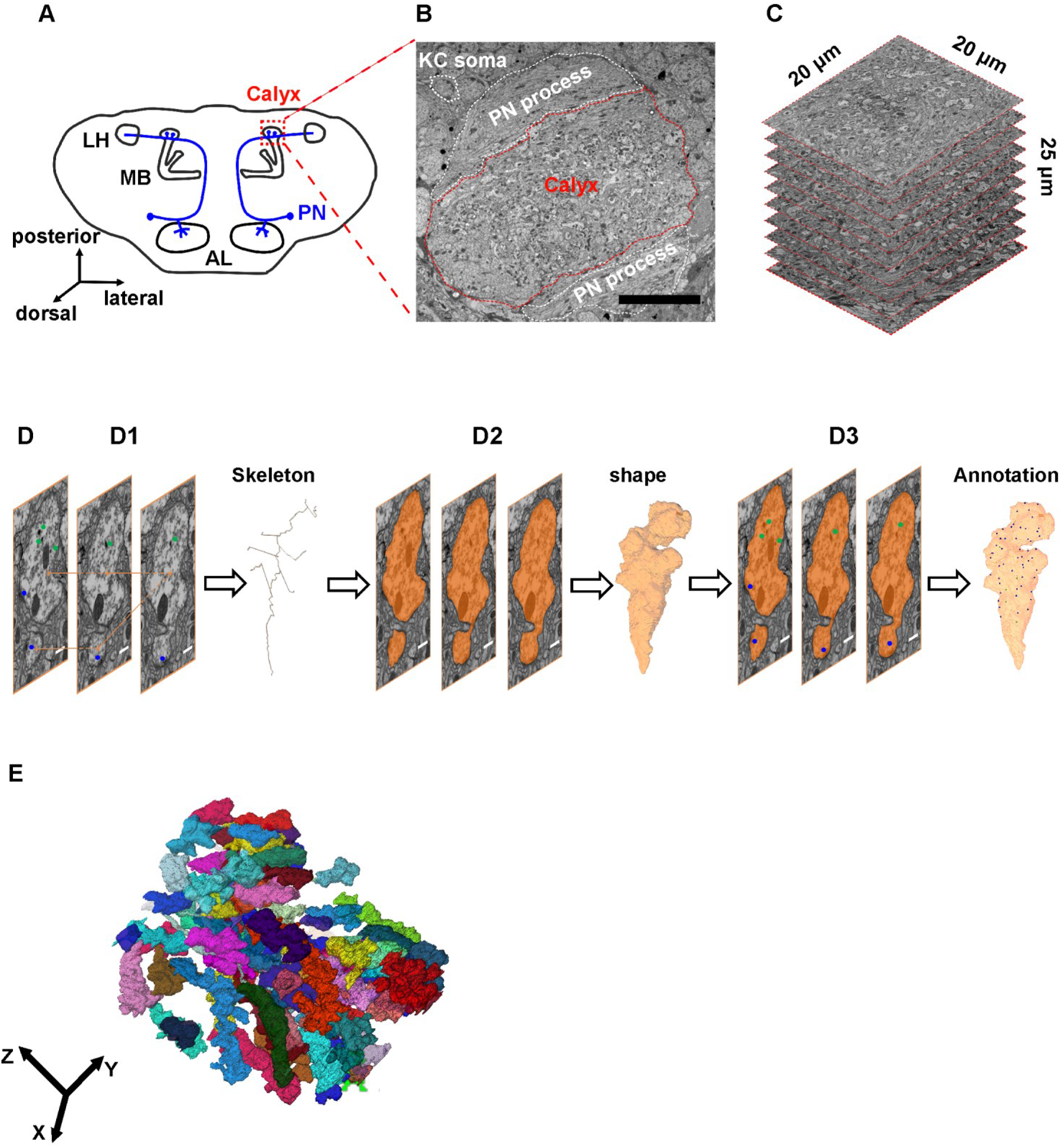
Volume reconstruction of projection neuron boutons in a reference region of the mushroom body calyx. (A) Schematic of olfactory projection neurons (PNs) in *Drosophila*. PNs connect the antenna lobe (AL) to the mushroom body (MB) and lateral horn (LH). The calyx is the input region of the MB. (B) Posterior view of the calyx under electron microscopy (EM). (C) Raw dataset from focused ion beam serial EM in the center of the calyx in a volume of 20 × 20 × 25 μm. (D) Workflow of volume reconstruction of PN axonal boutons. Scale bars = 5 μm. (E) Overall view of EM-reconstructed PN boutons.

PN boutons were identified in individual EM sections via four distinct features; large section area, enriched clear core vesicles, PAZs, and large internal mitochondria [13, 19]. The center of the same PN bouton was manually connected in consecutive sections to draw its skeleton (Figure 1 D1). The skeleton demonstrates the major axis and branches of a bouton. PN boutons were then volume reconstructed to depict their shapes. Based on their shapes, the three-dimensional features of each bouton such as volume and surface area were calculated (Figure 1 D2). Under EM most PAZs in insects such as *Drosophila* and *locusts* exhibit an electron dense ribbon called a T-bar, and DCVs are large electron-dense dots [5, 25]. Accordingly, all PAZs and DCVs in PN boutons were annotated (Figure 1 D3). To increase the accuracy of quantitative analysis of PAZs and DCVs, PN boutons that partially extended out of the EM dataset were not reconstructed (Figure S1A). Vesicles, mitochondria, and synapses are enriched in PN boutons, whereas PN fibers barely contain them (Figure S1B). This suggests that PN fibers may be involved in signal transduction in the neuron, but not communication between the neurons. Therefore, we focused on PN boutons, and PN fibers were not reconstructed. In total 89 PN boutons were reconstructed, and they were uniformly distributed in the EM dataset and exhibited variable size and shape. Most PN boutons were parallel with the z axis of the dataset (Figure 1E).

### Classification of PN boutons based on EM volume parameters

To investigate the distributions of DCVs and PAZs in PN boutons, we first examined the features and types of these PN boutons. Several morphological parameters of the reconstructed PN boutons were analyzed via Fiji-TrakEM2, and a PN bouton skeleton was generated. The skeleton provides several two-dimensional morphological features such as segment number and the total length of the skeleton (Figure 2A). Most PN boutons (80/89) contained 1–7 segments with a mean of approximately 5 (Figure S2A). The total length of PN bouton skeletons ranged from 2.3 μm to 53.9 μm, and the mean was 16.9 μm (Figure S2B). Total skeleton length was positively associated with segment number (Figure 2B), indicating that each segment has a similar length, and long PNs may contain a large number of segments.

**Figure 2:**
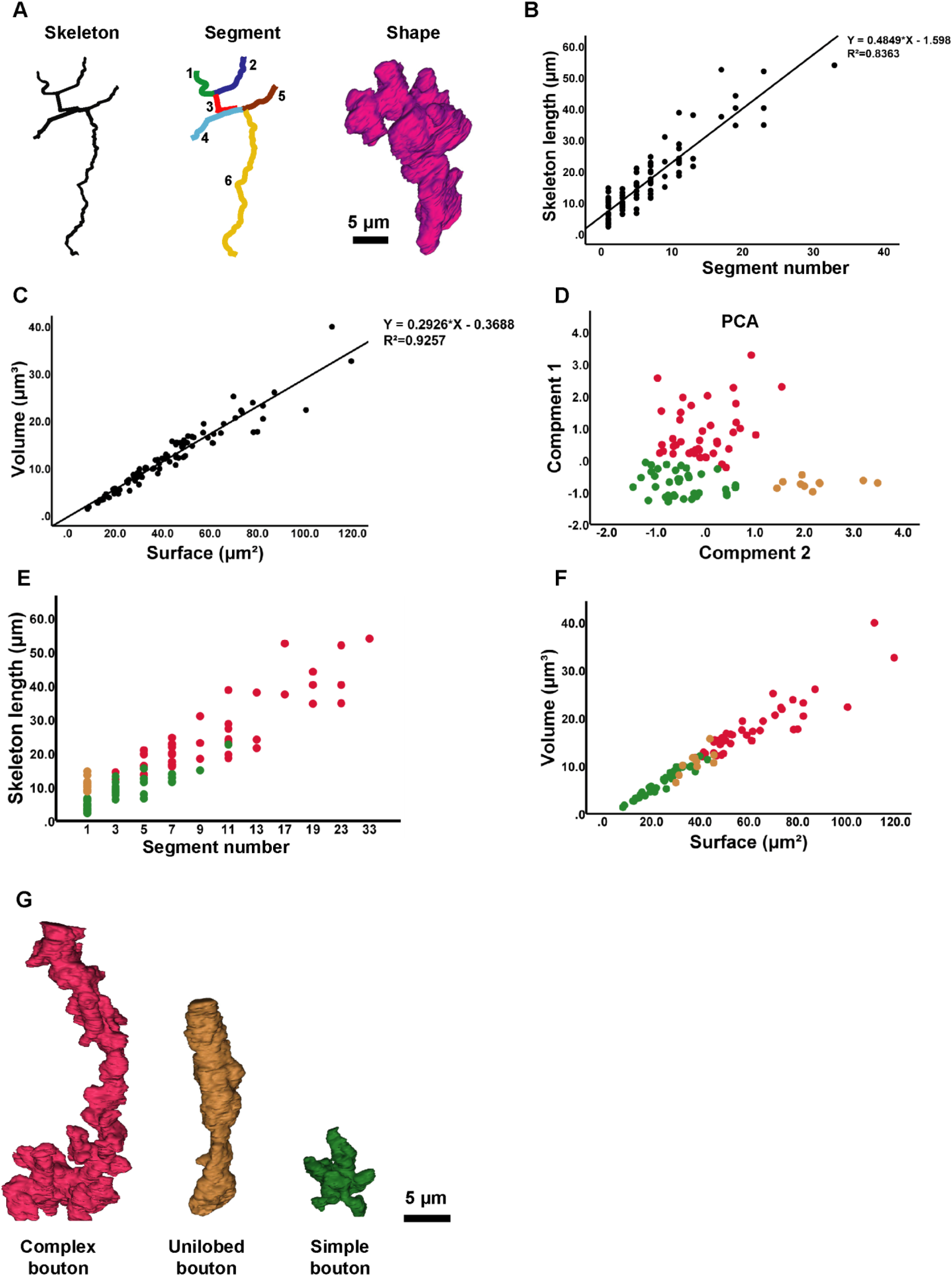
Morphological characteristics of PN boutons. (A) Segment and shape identification. Scale bars = 5 μm. (B) Skeleton length is positively correlated with segment number. (C) Bouton volume and surface area are positively correlated. (D) PN bouton classification based on morphological parameters. (E) Relationship between skeleton length and segment number in different types of PN boutons. (F) Relationships between volume and surface area in different types of PN boutons. (G) Schematic of three types of PN boutons. Scale bars = 5 μm.

The volume and surface area of PN boutons were then calculated to analyze three-dimensional features (Figure 2A). The surface area of most PN boutons was between 8.4 μm^2^ and 119.5 μm^2^ (mean 43.5 μm^2^), the volume of PN boutons ranged from 1.3 μm^3^ to 39.9 μm^3^ (mean 12.35 μm^3^), and 72/90 (80%) were between 10 μm^3^ and 30 μm^3^ (Figure S2). PN volume and PN surface area were positively correlated (Figure 2C), indicating that the surface area to volume ratio of PN boutons is similar regardless of their size.

Previous studies defined three types of PN boutons based on features determined via light microscopy (unilobed, clustered, and elongated), and two types based on features determined via electron microscopy (short segments and long slender axons) [13]. In the current study, after determining all the aforementioned morphological parameters all the reconstructed PN boutons were classified via automatic clustering. Principal component analysis was performed for total skeleton length, segment number, volume, and surface area, and two major components were extracted (Figure 2D). PN boutons were automatically clustered into three groups via Akaike information criterion-based two-step clustering (Figure 2D–G). Forty-two PN boutons exhibited long skeleton length and large segment number, surface area, and volume, and we named them complex boutons. Ten PN boutons exhibited just one segment with a long skeleton, and we named them unilobed boutons. Thirty-seven PN boutons exhibited short skeleton length, and small segment number, volume, and surface area, and we named them simple boutons (Figure 2E–G, S3).

### Ultrastructural distribution of PAZs and DCVs in PN boutons

In the central nervous system of *Drosophila*, most PAZs are electron dense ribbon T-bars, but some are non-ribbon type, and are attached by an adjunct postsynaptic density (Figure 3A–C). The neuromodulator-containing DCVs are electron-denser and larger than transmitter-contained clear core vesicles (Figure 3A–C). In accordance with the distinct criteria of PAZs and DCVs, we annotated all PAZs and DCVs in the volume reconstructed 89 PN boutons. At the subcellular level, as well as drifting in PN boutons, DCVs were often enriched in three regions; close to the membrane (Figure 3A), close to PAZ (Figure 3B), and close to mitochondria (Figure 3C). DCVs that are close to the non-PAZ membrane and PAZs may be respectively related to extra-synaptic and synaptic release of neuropeptide. There are few reports of DCVs close to mitochondria however, therefore their function is still unclear. PN boutons contained approximately 19 DCVs on average, with a maximum of 49 and a minimum of 1 (Figure S2E). There were approximately 44 PAZs in each PN bouton on average, with a maximum of 126 and a minimum of 9 (Figure S2F). To investigate the PAZ and DCV distributions in different types of PN boutons, we analyzed correlations between PAZ number, DCV number, and volume in individual PN boutons. PAZ number was positively associated with bouton volume, and DCV number was weakly associated with bouton volume (Figure 3D and 3E). These results indicate that PAZs are uniformly distributed in boutons, and DCV distribution is weakly associated with the morphology of PN boutons. PAZ and DCV numbers in three PN bouton types were investigated. Because complex, unilobed, and simple boutons have different volumes (Figure 2F), we predicted that there would be more PAZs in complex boutons, and no specific enrichment in different types of boutons. As predicted, there were significantly more PAZs in complex boutons than in unilobed boutons and simple boutons (Figure 3F). There were significantly more DCVs in complex boutons than in simple boutons, but the numbers of DCVs were similar in complex boutons and unilobed boutons (Figure 3G).

**Figure 3:**
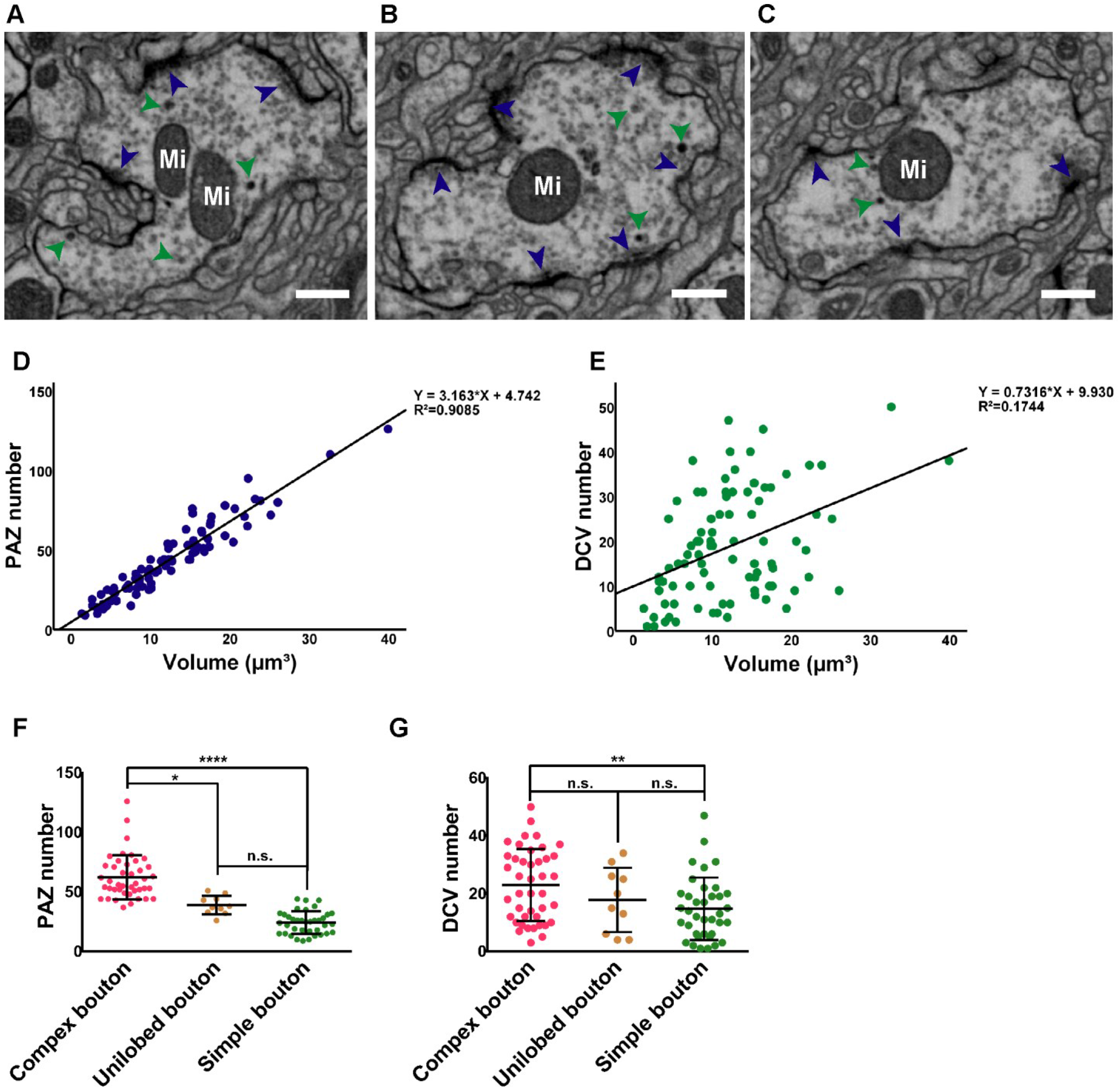
Distributions of presynaptic active zones and dense core vesicles in PN boutons. (A-C) Subcellular distributions of presynaptic active zones (PAZs; blue arrows) and dense core vesicles (DCVs; green arrows) in PN boutons. Most PAZs feature a T-bar ribbon (A and B), but a small number of PAZs have non-ribbon active zones (C). DCVs are varied in size and electron density (A and B). As well as drifting in boutons (A), DCVs are distributed in three patterns, close to the membrane (A), close to the T-bar (B), and close to mitochondria (Mi, C). Scale bar = 5 μm. (D) PAZ number was strongly linearly related to bouton volume (r^2^ = 0.91). (E) DCV number was not linearly related to bouton surface area (r^2^ = 0.17). (F) PAZ number in complex boutons was significantly larger than that in unilobed boutons and simple boutons. (G) DCV number in complex boutons was similar to that in unilobed boutons and larger than that in simple boutons (unpaired *t*-test). \**p* < 0.05, \*\**p* < 0.01, \*\*\**p* < 0.001, \*\*\*\**p* < 0.0001, n.s. = not significant

### DCVs are enriched in GH146-positive boutons

To investigate relationships between PAZ and DCV distributions, all of the reconstructed PN boutons were categorized into four clusters based on PAZ and DCV numbers via a Bayesian information criterion-based two-step clustering method (Figure 4A). PAZ and DCV numbers in different groups were compared to confirm the classifications (Figure 4B and 4C). Fewer boutons contained large numbers of both PAZs and DCVs, compared to other types (Figure 4D). RNAseq investigations indicated that PN subtypes labeled with abnormal chemosensory jump 6 (acj6) strongly expressed the sNPF gene, and PN subtypes labeled with ventral veins lacking (vvl) expressed the neuropeptide tachykinin gene [18]. acj6 and vvl were exclusively expressed in anterodorsal PNs and lateral PNs that were labeled with GH146-GAL4 [26, 27]. We then tested the hypothesis that the distribution of DCVs and PAZs may be associated with genetically different PN subtypes. To determine the DCV distribution in GH146-positive and GH146-negative PNs, GH146-positive boutons were labeled using UAS-mCD8:HRP which is driven by GH146-GAL4. The membranes of GH146-positive boutons were electron denser than those of GH146-negative boutons as determined via EM [28] (Figure 4E and 4F). More than half of GH146-positive PN boutons were in many-DCV clusters (clusters 1 and 2), whereas most GH146-negative boutons were in few-DCV clusters (clusters 3 and 4) (Figure 4G and 4H). These results indicate that as well as releasing neurotransmitters, GH146-positive PNs are also neuromodulators, but GH146-negative PNs may only release neurotransmitters to process olfactory information.

**Figure 4:**
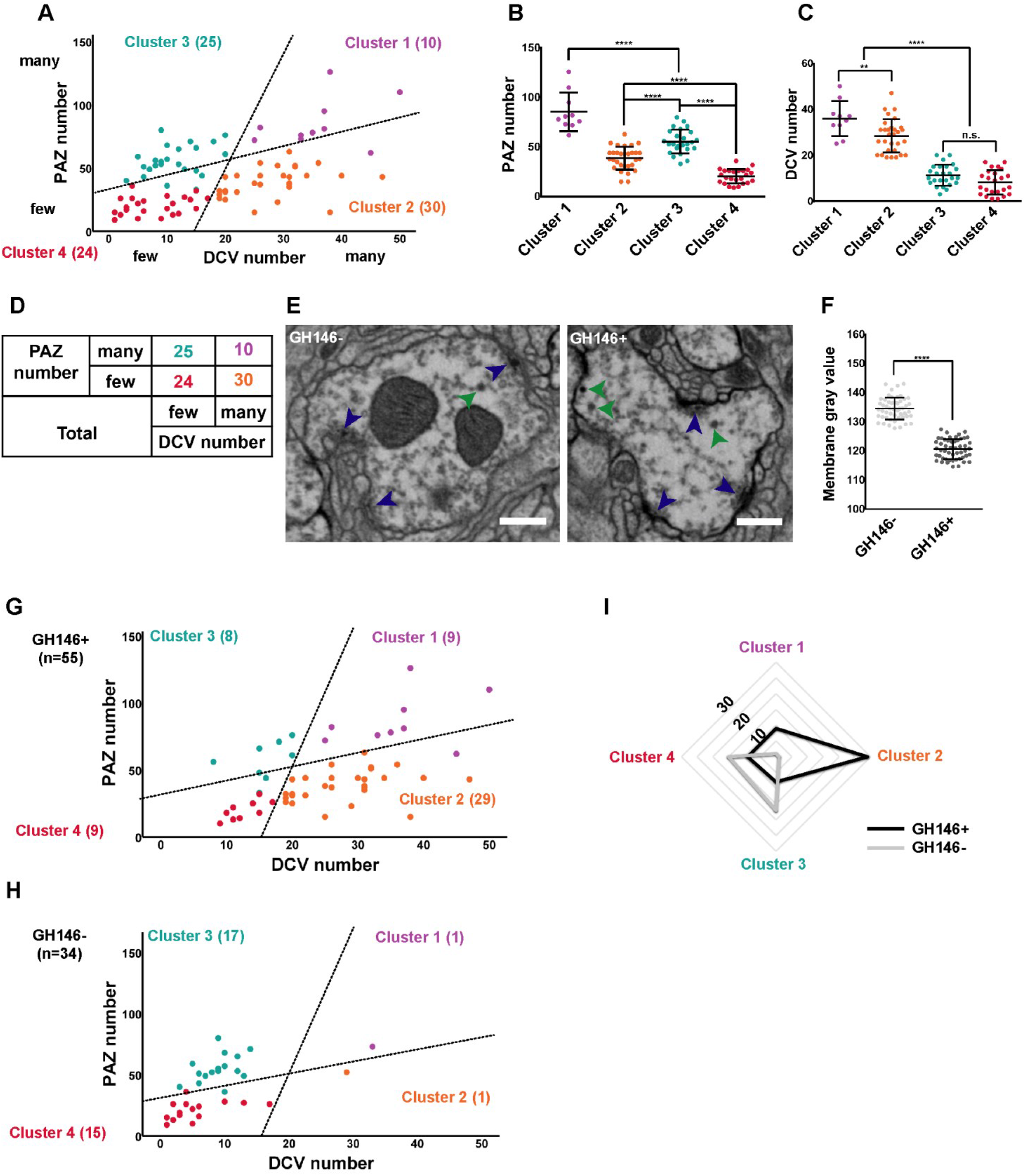
DCVs are enriched in GH146-positive PN boutons. (A) Clustering based on PAZ numbers and DCV numbers in individual PN boutons. Four clusters are evident; boutons containing many PAZs and many DCVs (violet), boutons containing many PAZs and few DCVs (cyan), boutons containing few PAZs and many DCVs (orange), and boutons containing few PAZs and few DCVs (red). (B and C) PAZ and DCV number comparisons in different clusters. (D) Bouton numbers in each cluster by PAZ and DCV number. (E) PAZs and DCVs in GH146-negative (GH146-) and GH146-positive (GH146+) boutons. Green arrows indicate DCVs and blue arrows indicate PAZs. Scale bar = 5 μm. (F) Membrane gray value in GH146-positive membranes was significantly larger than that in GH146-negative membranes. (G-I) Numbers of PAZs and DCVs in GH146-positive (G) and GH146-negative boutons (H). (I) Radar plots showing the number of GH146-positive boutons (black) and GH146-negative boutons (gray) in four clusters by PAZ and DCV number.

## Discussion

In the current study we volume-reconstructed 89 PN boutons in a reference region of the MB calyx in adult *D. melanogaster* and revealed some distribution features of DCVs and the neurotransmitter releasing sites, PAZs. Morphological parameter-based clustering indicated three different types of PN boutons in the calyx, including long and multiple-branched complex boutons, unilobed boutons with one long branch, and simple boutons with a few branches. At the subcellular level DCVs are located in four regions, including drifting in cytoplasm, and close to mitochondria, the T-bar, or the membrane. PAZs are uniformly distributed on the surface of all boutons at a density of 1 PAZ per square micron. GH146-positive boutons contained significantly more DCVs than GH146-negative boutons. This indicates that GH146-positive PNs release both neurotransmitters and neuromodulators, and GH146-negative PNs only release neurotransmitters. This study demonstrates the detailed distributions of PAZs and DCVs in different PN boutons, which will shed light on neuromodulator function in the olfactory circuit in *D. melanogaster*.

### Morphologically distinct PN bouton types

Olfactory PNs convey odor information from the antenna lobe to MB KCs by forming a microglomerulus, a structure where large PN boutons are surrounded by numerous tiny KC claws and some other MB extrinsic neuropiles, such as anterior paired lateral neurons and dopaminergic neurons [29–31]. Previous studies have shown that PN boutons in the MB calyx are distributed in a stereotypic pattern and exhibit no subregion preference [32]. Connections in the microglomerulus are the basis of divergent features for olfactory information transmission, and modulation by external neurons. Under light microscopy PN boutons reportedly exhibit three patterns; unilobed, elongated, and clustered [13]. In the present study, under EM PN boutons could be classified into three types based on their fine morphological features. Complex boutons have a large volume and a large surface area (Figure 2F and 2G) [13], therefore we propose that complex boutons are the same as clustered boutons. The unilobed boutons we identified had one long skeleton without branches (Figure 2E and 2G), therefore they are morphologically similar to elongated boutons under light microscopy. Unilobed boutons exhibit a short skeleton with small branches under light microscopy, as do simple boutons under EM. Axons undergo remodeling during development and in mature brains, including axon branching and pruning [33–35]. Thus, we propose that these three types of boutons may be mutually transformed, for example unilobed boutons may be the intermediate state between complex boutons and simple boutons, therefore their number is small.

### Distribution characteristics of PAZs in PN boutons

It is widely accepted that neurotransmitters are stored in clear core vesicles and are released into the synaptic cleft via PAZs [36, 37]. In *Drosophila* most PAZs feature an electron-dense ribbon called a T-bar, but there are a few PAZs without ribbons [13, 25]. Given the correlation between PAZs and clear core vesicles it is reasonable to surmise that boutons have the capacity to release neurotransmitters. The current study indicates that PAZ number is closely related to bouton volume, and that PAZs are uniformly distributed in PN boutons at a density of 1 PAZ per square micron. Therefore, the larger a bouton is, the more PAZs it contains. The results of the present study are consistent with a previous report suggesting that PAZ density is controlled in both neuromuscular junctions and the central nervous system [38]. Assuming that all PAZs have an equal propensity to release neurotransmitter, the neurotransmission capacity of PN boutons should correspond to their volume, therefore, larger boutons have higher neurotransmission capacity. In *Drosophila*, prolonged deprivation of synaptic transmission from PNs causes increased bouton size and PAZ density, as does aging [39, 40]. Unlike our definition of PAZ density—the number of PAZs per square micron—in the aforementioned studies PAZ density was calculated as the number of PAZs in individual boutons. We suggest that PAZ number per square micron is the key determinant of the efficiency of boutons with respect to information transmission, and it is controlled by unknown mechanisms [38]. Under disease conditions or in aged individuals, animals may compensate for reduced synapse release capacity by increasing bouton size and PAZ number, to maintain the transmission capacity of boutons. If that is true, what determines PAZ density and what changes PAZ density warrant further investigation.

### Characteristics of DCV distribution in PN boutons

In *Drosophila*, neuropeptides play key roles in the regulation of physiological and behavioral functions, including development, feeding, reproduction, and metabolism, and neuromodulation roles in olfaction, locomotor control, and learning and memory [41–48]. In the olfactory circuit, olfactory receptor neurons and MB KCs reportedly release neuropeptides that respectively facilitate and potentiate neurotransmission [9, 10]. Recently, single-cell sequencing results revealed that subtypes of PNs express neuropeptide gene [18], and the results of the current study constitute structural evidence that DCVs are enriched in GH146-positive PNs (Figure 4). It is thus reasonable to propose that PN subtypes labeled with GH146-GAL4 release neuropeptides as cotransmitters that modulate olfactory information transmission between PNs and KCs. As a typical system for studying the transformation from coarse coding to sparse coding, multiple components have been shown to underlie PN-KC transformation, including thresholding, input integration, sparse synaptic connection, and developmental plasticity [14–17]. Herein, we propose a new mechanism, *i.e*., that neuropeptides are involved in this process, because neuropeptides directly or indirectly modulate microcircuits [2]. Future functional studies will focus on investigating which neuropeptides are released by GH146-positive PNs, and how they are involved in PN-KC neurotransmission. Unlike previous studies in which a proportion of PN boutons reportedly contained no DCVs [13], in the present study all PN boutons contained DCVs, though a small number of PN boutons only contained a few (Figure 4D). This may be a result of some neuropeptides participating in axon guidance [49].

At the subcellular level neuropeptide is packaged into DCVs in endoplasmic reticulum, transported to axons via the trans-Golgi network, and released at synaptic or extra-synaptic sites [50]. The results of the current study indicate that DCVs are enriched in four regions, including drifting in cytoplasm, and close to PAZs and non-PAZ membranes (Figure 3A and 3B), and this constitutes ultrastructural evidence for DCV transportation in fly brain. Notably some DCVs in PN boutons were located close to mitochondria (Figure 3C) in the present study, and there are only a few reports derived from *C. elegans* indicating that neuropeptide release is related to mitochondria [51]. One possibility is that DCVs and mitochondria are transported together through the trans-Golgi network, and another possibility is that mitochondria-related DCVs are mitochondria-derived vesicles [50, 52].

In summary, in the current study HRP-labeled EM was used to reconstruct PN axonal boutons to reveal some new ultrastructural features of them with respect to bouton shape, PAZ distribution, and DCV distribution. These ultrastructural features constitute new evidence or cues for the study of PN function in olfactory circuits.

---

**Supplementary Figure 1:** Exclusive criteria for PN bouton tracing. (A) PN boutons that partially extended beyond the imaging field were excluded. (B) PN fiber that does not contain PAZs or DCVs, and was therefore excluded.

**Supplementary Figure 2:** Distributions of segment number, skeleton length, surface area, volume, DCV number, and PAZ number.

**Supplementary Figure 3:** Volume reconstruction of 89 PN boutons. Shapes of (A) 42 complex boutons, (B) 10 unilobed boutons, and (C) 37 simple boutons.

