## Supplementary materials for "Ultrastructural features of presynaptic active zones and dense core vesicles of olfactory projection neuron boutons in *Drosophila melanogaster*"

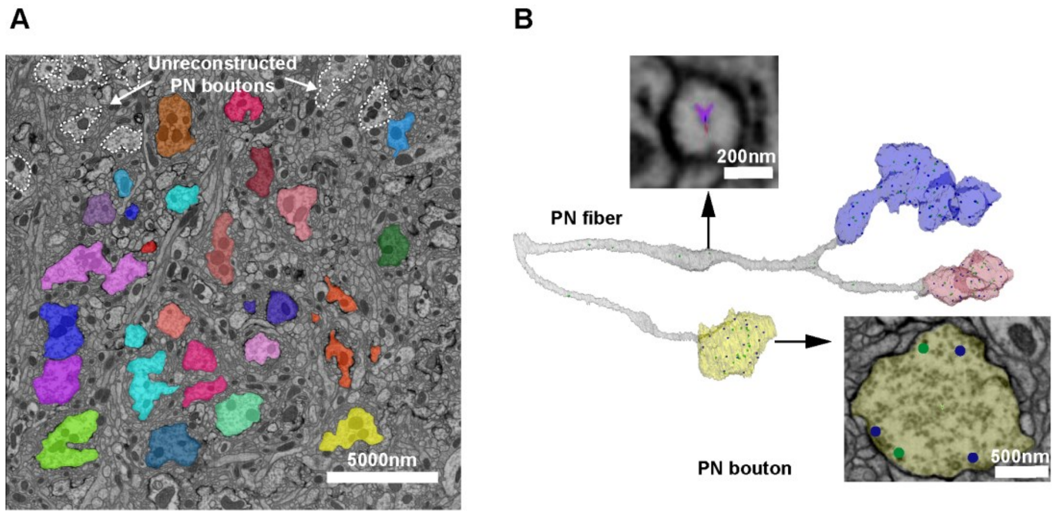


**Supplementary Figure 1:** Exclusive criteria for PN bouton tracing.

(A) PN boutons that partially extended beyond the imaging field were excluded. (B) PN fiber that does not contain PAZs or DCVs, and was therefore excluded.

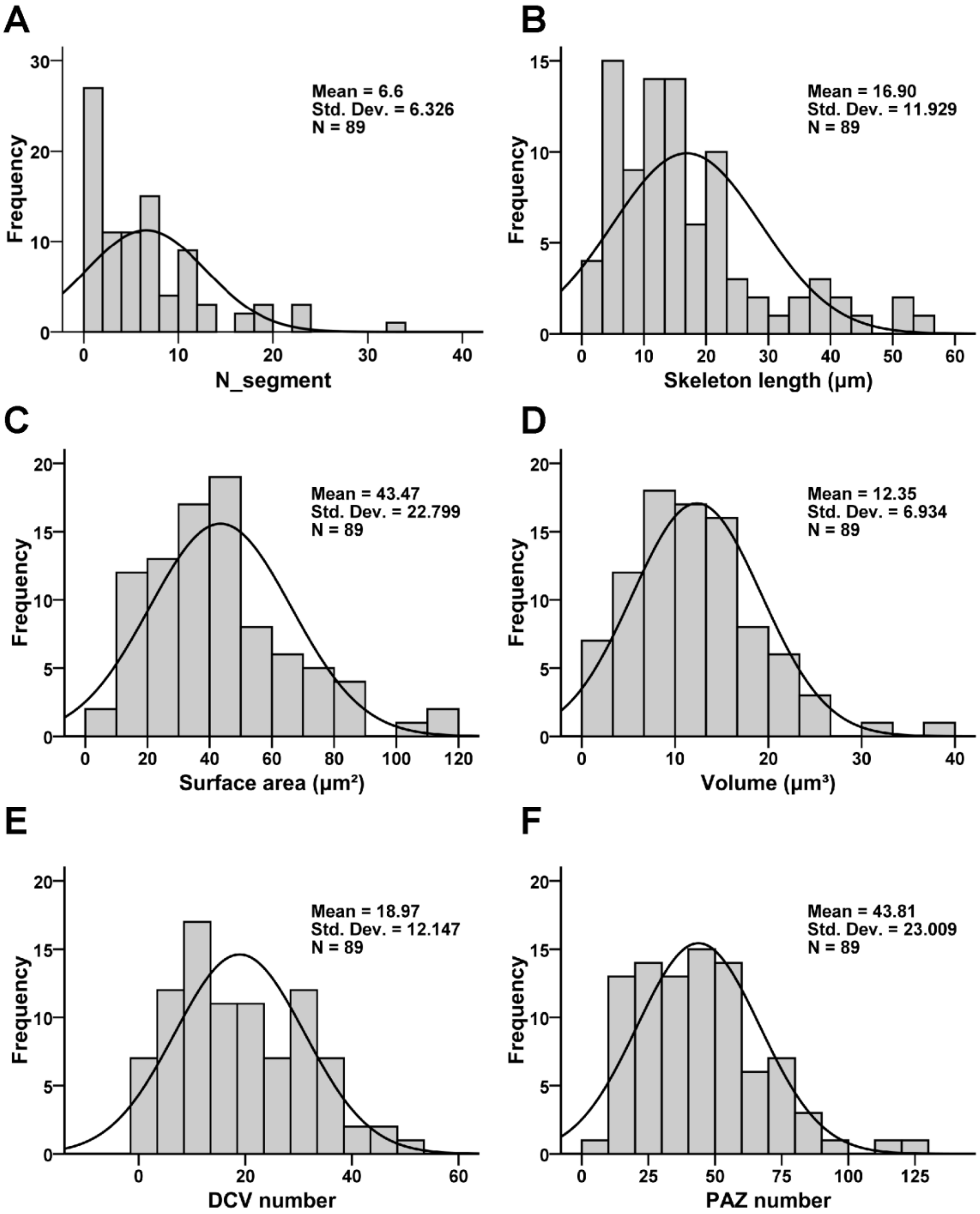


**Supplementary Figure 2:** Distributions of segment number, skeleton length, surface area, volume, DCV number, and PAZ number.

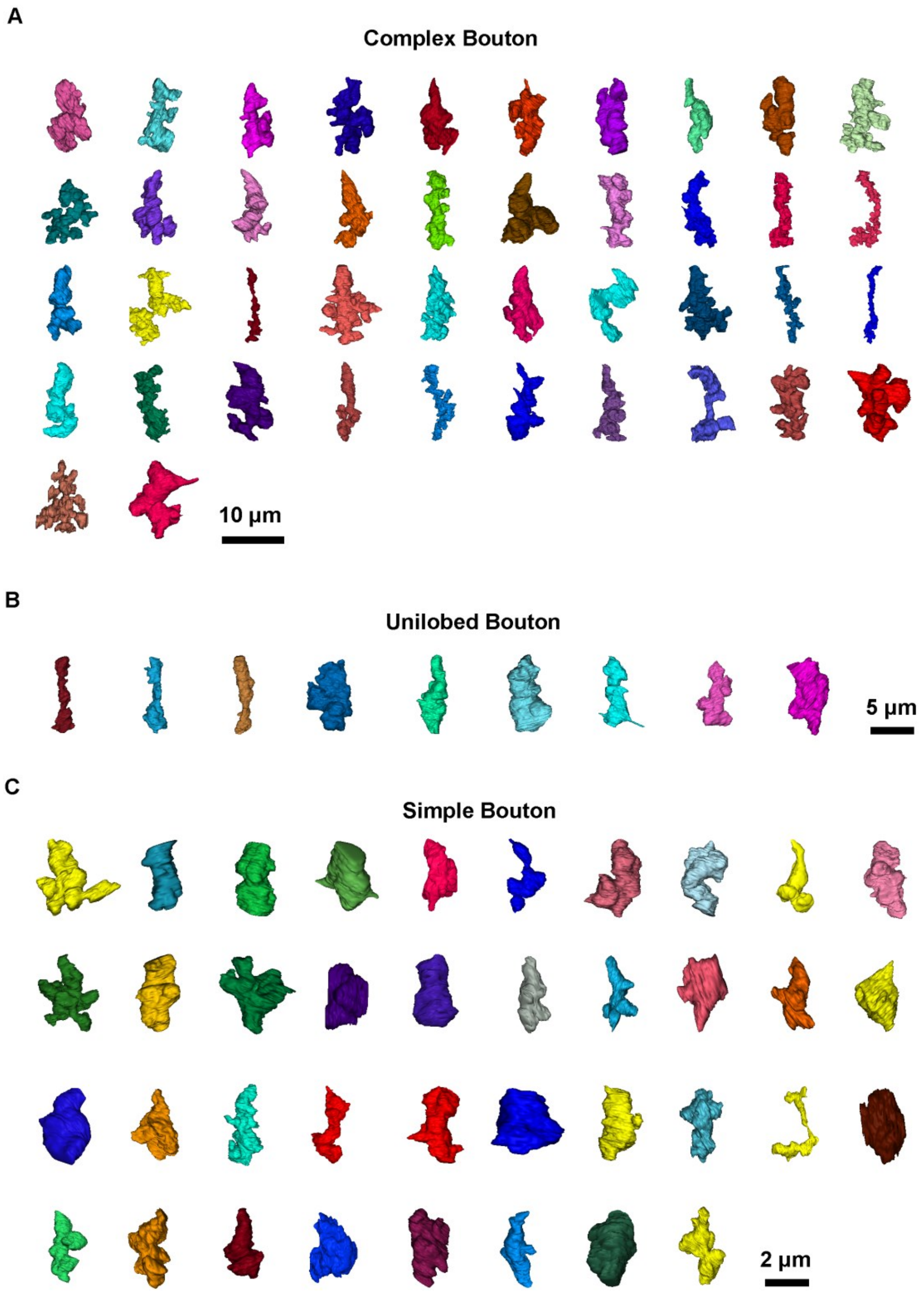


**Supplementary Figure 3:** Volume reconstruction of 89 PN boutons.

Shapes of (A) 42 complex boutons, (B) 10 unilobed boutons, and (C) 37 simple boutons.
